# The evolution of seasonal migration and the slow-fast continuum of life history in birds

**DOI:** 10.1101/2020.06.27.175539

**Authors:** Benjamin M. Winger, Teresa M. Pegan

## Abstract

Seasonal migration is a widespread adaptation in environments with predictable periods of resource abundance and scarcity. Migration is frequently associated with high mortality, suggesting that migratory species live on the “fast” end of the slow-fast continuum of life history. However, few interspecific comparative studies have tested this assumption and prior assessments have been complicated by environmental variation among breeding locations. We evaluate how migration distance influences the tradeoff between reproduction and survival in 45 species of mostly passerine birds that breed sympatrically in North American boreal forests but migrate to a diversity of environments and latitudes for the northern winter. We find, after accounting for mass and phylogeny, that longer distance migrations to increasingly amenable winter environments are correlated with reduced annual reproductive output, but also result in increased adult survival compared to shorter-distance migrations. Non-migratory boreal species have life history parameters more similar to long-distance migrants than to shorter-distance migrants. These results suggest that long-distance migration and other highly specialized strategies for survival in seasonal environments impose selection pressures that both confer and demand high adult survival rates. That is, owing to the reproductive cost of long-distance migration, this strategy can only persist if balanced by high adult survival. Our results reveal migratory distance as a fundamental life history parameter that predicts, and is inextricable from, the balance of survival and reproduction. Our study further provides evolutionary context for understanding the annual cycle demography of migratory species and the strategies long-distance migrants use to maximize survival on their journeys.

## Introduction

Annual fecundity and age-specific survival represent a fundamental tradeoff along the slow-fast continuum of life history (Stearns 1976; Reznick 1984; Saether 1988; Ricklefs 2000*a*; Martin 2002). Fecundity and survival operate in tension presumably owing to physiological constraints, though the evolutionary causes of this tradeoff remain debated (Gasser et al. 2000; Zera and Harshman 2001; Ricklefs and Wikelski 2002; Karagicheva et al. 2018). This tradeoff is integral to understanding the evolution of species’ behavioral and ecological attributes, because traits that promote survival may come at a cost to fecundity and vice versa (Clark and Martin 2007; Sol et al. 2016). However, the clarity of these patterns is often obfuscated by life history variation along ecogeographic gradients, necessitating hypothesis testing frameworks that distinguish between intrinsic constraints on life history and environmental variation in demographic parameters (Ricklefs 2000*a*; Ricklefs and Wikelski 2002). In this paper, we examine the evolutionary connections between seasonal migration—a widespread adaptation to seasonality in vagile animals—and the slow-fast continuum of life history. We show that the extraordinary long-distance migrations that characterize many temperate-breeding passerine birds are fundamentally shaped by a tradeoff between annual fecundity and adult survival. Further, we show that the migration distance of a species predicts its position along the slow-fast continuum—but in a direction counter to widespread assumptions about migration.

Seasonal migration is an annual round-trip between regions dedicated to reproduction and those dedicated to survival (Greenberg and Marra 2005), and as such is intrinsically connected to these two fundamental life history parameters. In birds, long a model for both life history theory (Ricklefs 2000*b*; Ricklefs and Wikelski 2002; Martin 2004) and migration research, a widespread view about migration and fecundity is that migration evolved in tropical species to increase reproductive output by exploiting seasonally available resources for breeding in the temperate zone while escaping competition in the tropics (Cox 1968, 1985). Migration has also been described as the costliest period of the annual cycle for migratory birds (Sillett and Holmes 2002). These ideas—that migration improves fecundity and is costly to survival—together have suggested (Bruderer and Salewski 2009) that the evolution of seasonal migration shifts migratory species towards the fast end of the slow-fast gradient (higher fecundity at a cost of lower survival; Sibly et al. 2012).

An alternative perspective is that migration evolves in seasonal areas to increase survival during the resource-depleted non-breeding season (Salewski and Bruderer 2007; Winger et al. 2019*a*). Under this view, migration does not evolve out of the tropics per se but rather evolves as a survival strategy in response to seasonality in a breeding range, regardless of the biogeographic origin of a lineage (Salewski and Bruderer 2007). That is, species exhibit migratory behavior in circumstances in which escape from the breeding grounds during a predictable period of resource scarcity improves annual survival and therefore the likelihood of achieving a breeding season the next year. This view casts migration as an adaptive strategy to seasonality analogous to hibernation, as opposed to an exploratory dispersal strategy to improve reproductive success relative to an ancestral tropical condition. If migration evolves to increase survival in the face of seasonality, then species that migrate to more favorable locations in the nonbreeding season might be expected to have higher survival than species that breed in the same seasonal environments but stay closer to their breeding grounds all year. If migration serves to promote annual survival, it should be expected to come at a cost to annual reproductive output (Ricklefs 2000*a*).

Despite the connections between life history theory and hypotheses for the evolution of migration in birds, the relationship between the migratory strategies employed by different species and tradeoffs in survival and fecundity has not been rigorously tested (Bruderer and Salewski 2009). The study of life history tradeoffs in birds has long focused on understanding ecogeographic patterns in reproductive strategies, such as larger clutch sizes in temperate versus tropical breeding birds (Ricklefs 2000*b*; Martin 2004) or the pace of growth and development across latitudes (Martin 2015). Owing to the general recognition of seasonal migration as a fundamental aspect of avian ecology and behavior, macroecological studies of global variation in avian reproductive output (Jetz et al. 2008; Sibly et al. 2012; Cooney et al. 2020) or survival (Muñoz et al. 2018; Bird et al. 2020) often include migratory status as a model covariate, but without an explicit hypothesis for its effect. These studies have often found migration to be associated with a “faster” life history strategy, such as high breeding productivity (Sibly et al. 2012). However, because migratory behavior is strongly correlated with seasonality and breeding latitude, migratory species are likely to exhibit life history conditions associated with breeding at high latitudes, such as relatively large clutch sizes (Jetz et al. 2008). Thus, a direct influence of migratory behavior on the slow-fast continuum may not be revealed by global analyses whose primary axis of variation is latitudinal.

A second area of relevant research has focused on seasonal variation in survival of migratory birds to improve full annual cycle population models (Faaborg et al. 2010; Hostetler et al. 2015; Marra et al. 2015). In a highly cited landmark study, Sillett and Holmes (2002) estimated the mortality associated with migration in a boreal-breeding Neotropical migratory songbird, the Black-throated Blue Warbler (*Setophaga caerulescens*). They found that most mortality in this species occurred during migration as opposed to the breeding (summer) or non-breeding (winter) “stationary” periods (Sillett and Holmes 2002). Although the goals of this and similar studies have been to compare seasonal variation in survival rates of individual migratory species, their results have sometimes been generalized to a paradigm that migration, broadly, is a strategy that carries a high survival cost (Newton 2007; Faaborg et al. 2010; Sibly et al. 2012; Somveille et al. 2019). Yet, it has more rarely been asked how the survival costs associated with long-distance migration compare to an alternative strategy of staying closer to the breeding grounds year-round (Greenberg 1980; Sherry and Holmes 1995; Bruderer and Salewski 2009). Though migration clearly requires substantial energy and presents a risk to survival, so does maintaining homeothermy and body condition in regions with cold temperatures or scarce food resources (Ricklefs 1980; Swanson and Garland 2009). Further, recent studies in other migratory species, especially non-passerines, have discovered high rates of adult survival associated with long-distance migration, calling into question the generality of a high mortality cost of migration (Lok et al. 2015; Conklin et al. 2017; Senner et al. 2019).

The consequences of migration for tradeoffs between survival and reproduction have been explored somewhat in studies of intraspecific variation in migratory behavior (Ketterson and Nolan 1982; Alves et al. 2013; Ely and Meixell 2015; Lok et al. 2017; Zúñiga et al. 2017; Buchan et al. 2019), but less so in the context of understanding the evolution of species-level life history tradeoffs and hence the position of species along the slow-fast continuum. Greenberg (1980) proposed the time allocation hypothesis to explain the relationship between migration, survival and reproduction. This study suggested that migratory species invest less time in breeding than temperate non-migratory species but compensate for lower fecundity by investing in migrations to locales with resources that support increased winter survival. A small number of studies have subsequently demonstrated that temperate-breeding migratory species have lower annual fecundity than temperate residents or short-distance migrants (Mönkkönen 1992; Böhning-Gaese et al. 2000; Bruderer and Salewski 2009). Research on other taxa has also found that migration distance may be positively, and counterintuitively, correlated with adult survival rates (Conklin et al. 2017) or longevity (Møller 2007). However, most previous interspecific comparisons have tested the relationship between migratory behavior and either survival and reproduction in isolation—rather than the tradeoffs among these parameters—and have often not been able to control for environmental heterogeneity among breeding locations that could influence demographic parameters (Ricklefs 2000*a*).

Here, we assess the influence of seasonal migration on the slow-fast continuum of life history in a geographically and ecologically circumscribed system: small-bodied, mostly passerine birds breeding in the boreal forests of eastern and central North America (Fig. 1). This system is well-suited to assessing the impact of migratory behavior on life history because of the high diversity of similarly-sized species that breed at the same latitudes and in the same habitat but spend the nonbreeding season in drastically different regions (from year-round residents that do not leave the frozen boreal region in the winter to long-distance migrants that migrate to South America; Fig. 1). By integrating multiple published long-term datasets on population demographic parameters from the boreal region, we test how variation in migratory distance influences species’ life history. First, we test the time allocation hypothesis (Greenberg 1980; Bruderer and Salewski 2009) that long-distance migratory species invest less time during the annual cycle in reproduction than co-distributed short-distance migrants or residents. Second, we test whether increased migratory distance and decreased time allocated to reproduction is associated with reduced annual fecundity. Third, we assess the influence of migration distance on annual adult survival and predict that if longer-distance migrations reduce fecundity, these migrations should also be associated with increased annual survival.

**Figure 1.**
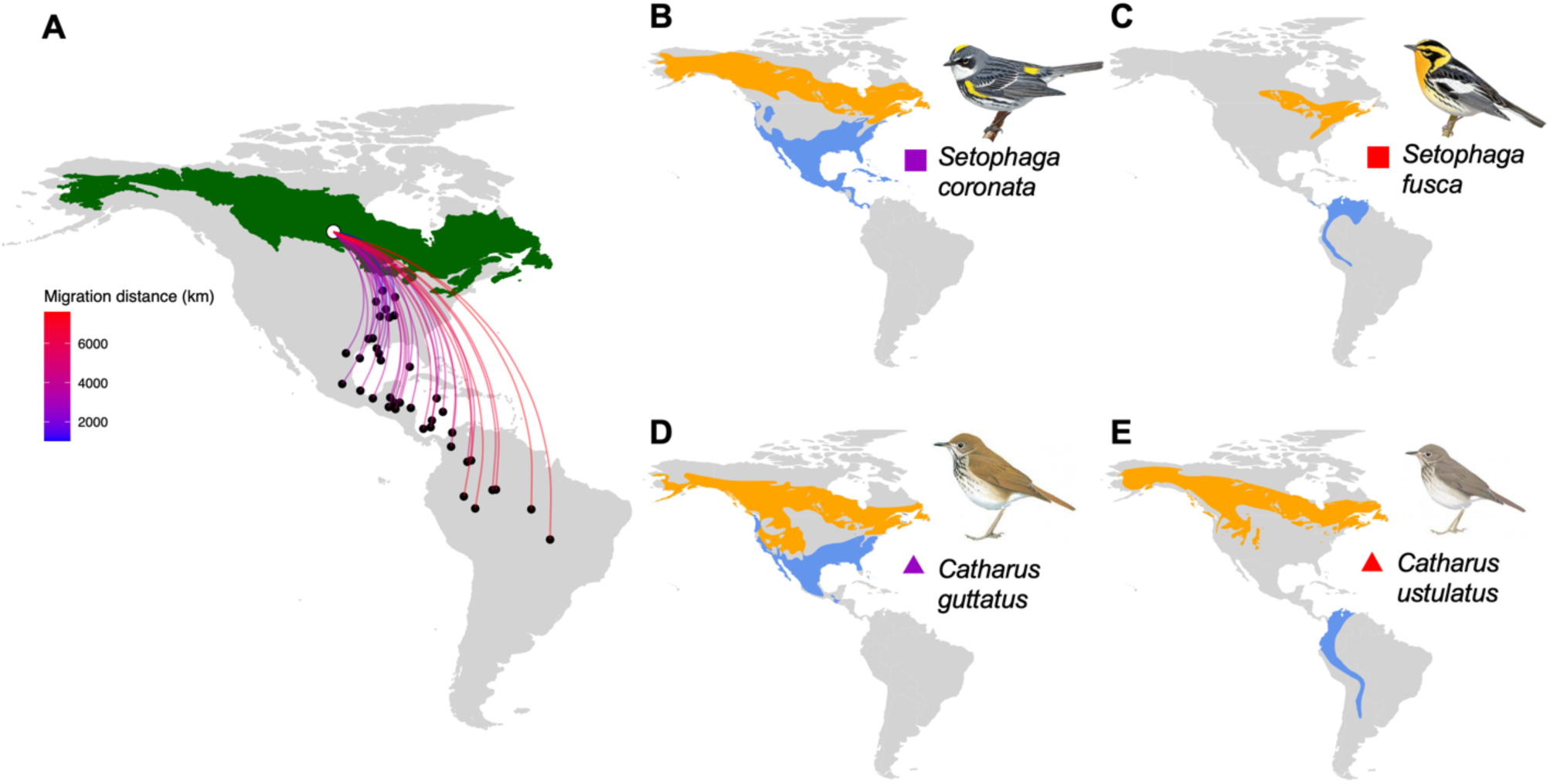
A) The 45 species in our study (Table S1) broadly co-occur during the breeding season across the boreal forest belt (green, indicating ecoregions “northern forests”, “taiga” and “Hudson plain”; Omernik and Griffith 2014) but winter in disparate locations. Hypothetical migration routes are depicted between a single breeding location and the centroid of the winter ranges of each of 39 migratory species (Methods). B-E) Example species representing short- (B, D) and long- (C, E) distance migratory species in pairs in each of two genera. These example species are highlighted in Figs. 2-4 with the squares (*Setophaga)* and triangles (*Catharus*) colored according to migratory distance in panel A. Species maps from BirdLife International and Naturserve (2014). Illustrations reproduced by permission of Lynx Edicions.

If migratory distance away from boreal breeding grounds influences survival, this relationship may be mediated by disparate conditions experienced on the wintering grounds. Are the costs of long-distance migration offset by benefits of more distant wintering grounds (Somveille et al. 2018)? To address this question, we test whether interspecific variation in migratory distances is correlated with macroecological proxies for winter environment (temperature, precipitation and primary productivity) and whether these variables influence annual survival. Survival could also be influenced by the climatic similarity of migratory species’ breeding and winter ranges during their summer and winter, respectively, which affects the breadth of conditions to which they must adapt throughout the year. Winter ranges of migratory birds tend to occur in areas that are climatically more similar to the breeding grounds than expected by chance, suggesting that “niche tracking” underlies migratory movements at macroecological scales (Gómez et al. 2016; Zurell et al. 2018; Somveille et al. 2019). To test whether niche tracking provides a survival benefit, we test how niche overlap between the breeding and winter ranges (Zurell et al. 2018) is correlated with migration distance and whether it predicts annual survival.

## Methods

### Study system

Our study system comprises 39 migratory and 6 non-migratory species of birds (41 passerine species from 11 families and 4 woodpecker species) breeding in forested habitats in the boreal forest belt of eastern and central North America (Fig. 1, Table S1). Together, these species comprise a regional community of breeding birds in the northern spring and summer but they spend the northern winter in disparate locales. Because body size is an important axis of the slow-fast continuum (Ricklefs 2000*a*; Martin 2004), we focused on small-bodied birds that can reproduce beginning at ∼1 year of age. Most species ranged from 6-33 grams (median 11 grams), but 7 species were larger (50-87 g). In exploratory analyses, we found similar results when restricting analyses to the set of species < 33 grams.

We included species that breed in bog, spruce-fir and mixed forested habitats throughout the boreal and hemiboreal region (Fig 1). To control for latitude, we excluded species primarily restricted to more northern treeline habitats of the taiga (e. g., *Catharus minimus, Setophaga striata*) and those primarily restricted to more southerly latitude (e. g., *Piranga olivacea, Setophaga pinus*) even if their breeding ranges overlapped marginally with our region. We included species whose ranges extend beyond the boreal region if the species breeds broadly throughout the boreal forest (e.g., *Picoides villosus, Vireo olivaceus*) and had available data from boreal populations. We included species with regular seasonal migrations and those known to be primarily non-migratory but excluded species that primarily undergo facultative irruptive or nomadic movements (e.g., *Sitta canadensis, Loxia leucoptera*) due to the difficulty of defining migratory patterns or breeding periods.

### Live history predictors: mass and migration

Mass data were obtained from Dunning (2008) or Billerman et al. (2020). We used a binary predictor, migratory status, to indicate whether or not a species engages in regular seasonal migration. We estimated migration distance as the geodesic distance between the centroids of the breeding and wintering range of each species, including any year-round portions in both the breeding and wintering range when calculating centroids (BirdLife International and Naturserve 2014). To ensure that our migration distance estimate was as accurate as possible for populations breeding in the boreal forest belt, we used only the portions of breeding ranges overlapping with boreal forest ecoregions (Omernik and Griffith 2014; Level I ecoregions “northern forests,” “taiga,” and “Hudson plain”; Fig. 1) in the calculation of breeding range centroid. We also excluded any portions of wintering ranges within or west of the Rocky Mountains because these areas are generally used by more western breeding populations (Kelly and Hutto 2005; Kardynal and Hobson 2017). Geographic calculations were made using geospatial packages in R (Bivand and Rundel 2019; Bivand et al. 2019; Hijmans 2019).

### Life history outcome variables

We gathered published data related to time allocation for breeding, developmental duration, annual reproductive output and annual adult survival (Table 1). The data we compiled was collected mainly within the boreal region (Fig. 1) to control for latitudinal variation in demographic parameters.

**Table 1:** Description of outcome variables. Summary of life history variables used in study and their data sources. See Methods for further details.

| Variable | Description | Source |
| --- | --- | --- |
| **Time Allocation** |  |  |
| Intermigratory Period | Interval between peak median spring and fall migration through Chicago, IL | (Sullivan et al. 2009) |
| Egg Interval | Breadth of dates that nests with eggs have been found in Ontario; represents breadth of the breeding cycle for a species in the boreal region. | (Peck and James 1983, 1987) |
| **Developmental Duration** |  |  |
| Incubation Period | Average of individual incubation periods in Ontario; represents individual-level embryonic developmental rate. | (Peck and James 1983, 1987) |
| **Reproduction** |  |  |
| Clutch Size | Average of clutch size ranges from Ontario | (Peck and James 1983, 1987) |
| Fecundity | Average clutch size x maximum number of successful broods per year | (Peck and James 1983, 1987; Billerman et al. 2020) |
| Reproductive Index | Estimated fledglings per adult from long-term capture-mark-recapture study in boreal region | (Desante et al. 2015) |
| **Survival** |  |  |
| Annual Adult Survival ($\phi$) | Apparent annual adult survival ($\phi$) from long-term capture-mark-recapture study in boreal region | (Desante et al. 2015) |

We estimated the number of days each species invests in breeding. Comparable data on the full breeding cycle (from establishment of territories and building of nests through fledging of chicks) was not available for the boreal region for many species, so we used three proxies.

First, we estimated the amount of time each species spends on or near its breeding grounds by using eBird data (Sullivan et al. 2009) to calculate the interval of time between spring and fall migratory passage through Chicago, IL, USA. Migratory passage through Chicago is a reasonable proxy for time spent on the breeding grounds for the migratory species in this study because Chicago is near to but not within the boreal region (approximately 350 km to the southern edge of the boreal forest), it is a stopover location common to all the species, it experiences a high density of bird migration, and an active community of birdwatchers generate ample eBird data (Winger et al. 2019*b*). We analyzed eBird data from Cook County, IL from 2000-2017. For each species in each year and season (spring and fall), we calculated the date with the most eBird records for that species and considered this to be the date of “peak migration”. We calculated the average number of days between the median peak spring and median peak fall migration dates across all years for each species. This “intermigratory period” serves as a proxy for the relative amount of time species spend on or near their breeding grounds.

Second, we gathered data specific to the boreal region on the egg laying period. We used data from *Breeding Birds of Ontario: Nidiology and Distribution* (Peck and James 1983, 1987), henceforth “*Nidiology*.” *Nidiology* summarizes nearly 85,000 nest records of birds breeding in Ontario, Canada, collected mostly by the Ontario Nest Records Scheme initiated in 1956. We used this volume because it is more specific to boreal latitudes and more consistent in information than more general sources on breeding birds (Billerman et al. 2020). We used *Nidiology’s* records of the earliest and latest egg dates to calculate the number of days over which each species lays or incubates eggs (“egg interval”; Table 1).

Third, we used *Nidiology* to calculate the average individual incubation period for each species (“incubation period”; Table 1). Egg interval represents the temporal breadth of the breeding cycle for a species in the boreal region, whereas incubation period represents an individual-level and embryonic developmental duration.

We calculated average clutch size from the range of clutch sizes listed in *Nidiology* for each species. We calculated annual fecundity as (average clutch size) x (maximum number of successful broods) per season. *Nidiology* did not contain information on number of broods, so we referred to Billerman et al. (2020) for information on number of broods. Data on incidence of double brooding were not available; we marked species as double brooders if species accounts indicated that an additional brood may be raised following a successful first brood.

We also compiled data on reproductive output from the Vital Rates of North American Birds project (henceforth, *Vital Rates;* Desante et al. 2015) for 29 species with available data. *Vital Rates* provides estimates of demographic parameters derived from widely implemented constant-effort capture-mark-recapture surveys from 1992-2006, as part of the Monitoring Avian Productivity and Survivorship (MAPS) program. We used *Vital Rate’s* Reproductive Index, an estimate of the number of young birds produced per adult annually, calculated using effort-corrected generalized linear mixed models (Desante et al. 2015). *Vital Rates* presents estimates of Reproductive Index from different geographic regions using Bird Conservation Regions (Bird Studies Canada and NABCI 2014) to delineate boundaries for study sites according to biomes. We calculated mean Reproductive Index for our species by prioritizing BCRs 8 and 12, which broadly overlap with eastern and central boreal forests, and substituting mean estimates BCRs 6, 7, 13, or 14 when necessary. We did not include any parameters that *Vital Rates* flagged as not usable due to unreliable model estimates.

We similarly estimated annual adult survival from boreal-specific regions using *Vital Rates’* Adult Apparent Survival Probability (*ϕ*, Cormack-Jolly-Seber estimation), which is “an estimate of the annual probability that a resident bird that was alive and present at the station in year t will also be alive and present in year t+1” (Desante et al. 2015).

### Modeling

We used linear models to test hypotheses on the relationship between seasonal migration and life history. For clutch size, we used average clutch sizes, meaning that these data were not integer counts. We modeled the relationship between the intermigratory period (of migratory species only) and migration distance, and between egg interval and both migratory status and migration distance. For the 5 life history outcome variables related to developmental duration, reproductive output and survival (Table 1) we also assessed the influence of body size. For these 5 variables we fit three models: One model with mass as a predictor, a second model with migration status and migration distance as predictors, and a third model with all three predictors. We log-transformed mass and the outcome variables because many biological processes scale non-linearly with body size, and centered and standardized migration distance. For each outcome variable, we compared the three models with the second-order Akaike Information Criterion (AICc) using *MuMIn* (Bartón 2019) to assess the relative performance of migration and mass as predictors. Models for different outcome variables contain slightly different subsets of species based on the availability of data on the outcome variable (Tables 1, S1).

We tested whether the data were best modeled using Phylogenetic Generalized Least Squares (PGLS) or Ordinary Least Squares (OLS) by fitting an OLS full model (using all relevant predictors) and then using the function phylosig from phytools (Revell 2012) to test for phylogenetic signal (λ) in the model’s residuals (Revell 2010). Although controlling for shared phylogenetic ancestry is important in comparative analyses, PGLS has the potential to reduce model accuracy when a model’s residuals do not have phylogenetic signal (Revell 2010). For a phylogeny, we built a consensus tree with data from birdtree.org (Jetz et al. 2012), using procedures described in Pegan and Winger (2020). For models with significant λ, we performed PGLS modeling with the gls function in *nlme* (Pinheiro et al. 2019), including a correlation structure of expected phylogenetic covariance among species according to a Brownian motion model (function “corBrownian” in *ape*; Paradis and Schliep 2019). For response variables that did not have significant phylogenetic signal in model residuals, we used OLS. We also present PGLS analyses of all outcome variables (Table S3) for which we jointly estimated Pagel’s λ and the model using “corPagel” in *ape* (Revell 2010; Paradis and Schliep 2019).

### Winter climate and niche tracking

To test whether differences in annual survival across species with different migratory distances are associated with differences in winter conditions, we assessed winter climate and primary productivity across the winter ranges of each species. We used eBird (Sullivan et al. 2009) records for each species from November to February (all years) and retained all points that fell within a species’ typical winter range (BirdLife International and Naturserve 2014) to exclude extralimital vagrants. We excluded *Catharus fuscescens* and *Oporonis agilis* due to a paucity of winter eBird records. We downloaded month-level climate data (Fick and Hijmans 2017) at 30s resolution and NDVI data (Pinzon and Tucker 2014), averaged from the years 2000-2010. We filtered rasters to contain one eBird record per grid cell to mitigate spatial bias and used these points to calculate species-level means for winter temperature, precipitation, and normalized difference vegetation index (NDVI). We then conducted a principal components analysis to reduce temperature, precipitation, and NDVI to a single species-level value (PC1, representing winter climate) and assessed the correlation between migratory distance and winter climate PC1.

We also assessed the relationship between migratory distance and an estimate of the overlap of breeding and non-breeding season climatic niches from Zurell et al. (2018). Niche overlap represents the similarity between the conditions (climate and NDVI) encountered in the breeding range during breeding months, and the winter range during winter months. Finally, we analyzed the effect of winter climate PC1 and niche overlap on annual adult survival (*ϕ*) in separate models that included mass as a covariate, after log-transforming *ϕ* and mass.

## Results

In most cases, models with migration variables (distance or status) had lower AICc scores (Table S2) than models containing only mass as a predictor, and migration variables were significantly (p<0.05) associated with most life history outcome variables after controlling for mass (see Table 2 for details), indicating that migration explains mass-specific variation in each outcome variable.

**Table 2:** Results of best-fit models. Summary of best-fit models for each life history variable. Variables without coefficients were not included in the best-fit model (see Table S2 for AICc model comparison results). Coefficients with p < 0.05 are italicized and bold and considered statistically significant in the text. Migration distance was centered and standardized. Mass was log-transformed and outcome variables whose best model included mass were also log-transformed. “Method” notes whether the model type was OLS or PGLS, based on significance of λ at p < 0.05 in a test of phylogenetic signal of model residuals (see Methods). Column “n” shows the number of species used in each model, based on data availability (Table S1). Across 39 migratory species, migration distance shows a significant negative relationship with breeding season length (intermigratory period and egg interval), clutch size and fecundity but a significant positive relationship with annual adult survival. By contrast, the 6 non-migratory species have a shorter egg interval, lower fecundity and higher survival (Fig. 3).

| Response | Mass | Migratory Status<br>(Nonmigratory) | Migration Distance | Method | $\lambda$ | n |
| --- | --- | --- | --- | --- | --- | --- |
| **Time Allocation** |  |  |  |  |  |  |
| Intermigratory Period |  |  | **-26.11** | OLS | 0.68 | 39 |
| Egg Interval |  | **-30.14** | **-9.09** | OLS | 0.47 | 41 |
| **Developmental Duration** |  |  |  |  |  |  |
| Incubation Period | 0.07 |  |  | PGLS | **0.73** | 31 |
| **Reproduction** |  |  |  |  |  |  |
| Clutch Size | **-0.17** | -0.07 | **-0.11** | OLS | 0.45 | 43 |
| Fecundity | **-0.16** | **-0.62** | **-0.33** | OLS | 0.21 | 43 |
| Reproductive Index |  | 0.33 | -0.09 | OLS | 0 | 29 |
| **Survival** |  |  |  |  |  |  |
| Annual Adult Survival ($\phi$) | **0.42** | **0.66** | **0.18** | PGLS | **1.03** | 28 |

For two life history outcome variables (incubation period and annual adult survival), we highlight results of phylogenetic generalized least squares regression (PGLS). For the remaining variables, we present the results of OLS regression because we found that PGLS was not justified based on lack of phylogenetic signal in the model residuals (Methods).

### Migration distance and life history

Among the migratory species in our study, the variables representing time allocation in breeding were significantly negatively associated with migration distance (Table 2). The analysis of intermigratory period indicates that long distance migratory species arrive later and spend significantly less time on or near their breeding grounds before departing earlier for fall migration than short-distance migratory species (Table 2, Fig. 2). For an increase in migration distance of one standard deviation, or about 2000 km, our best model predicts a decrease in intermigratory period of about 26 days. This pattern was corroborated by egg interval, which predicted an egg laying period about 9 days shorter per standard deviation increase in migration distance.

**Figure 2.**
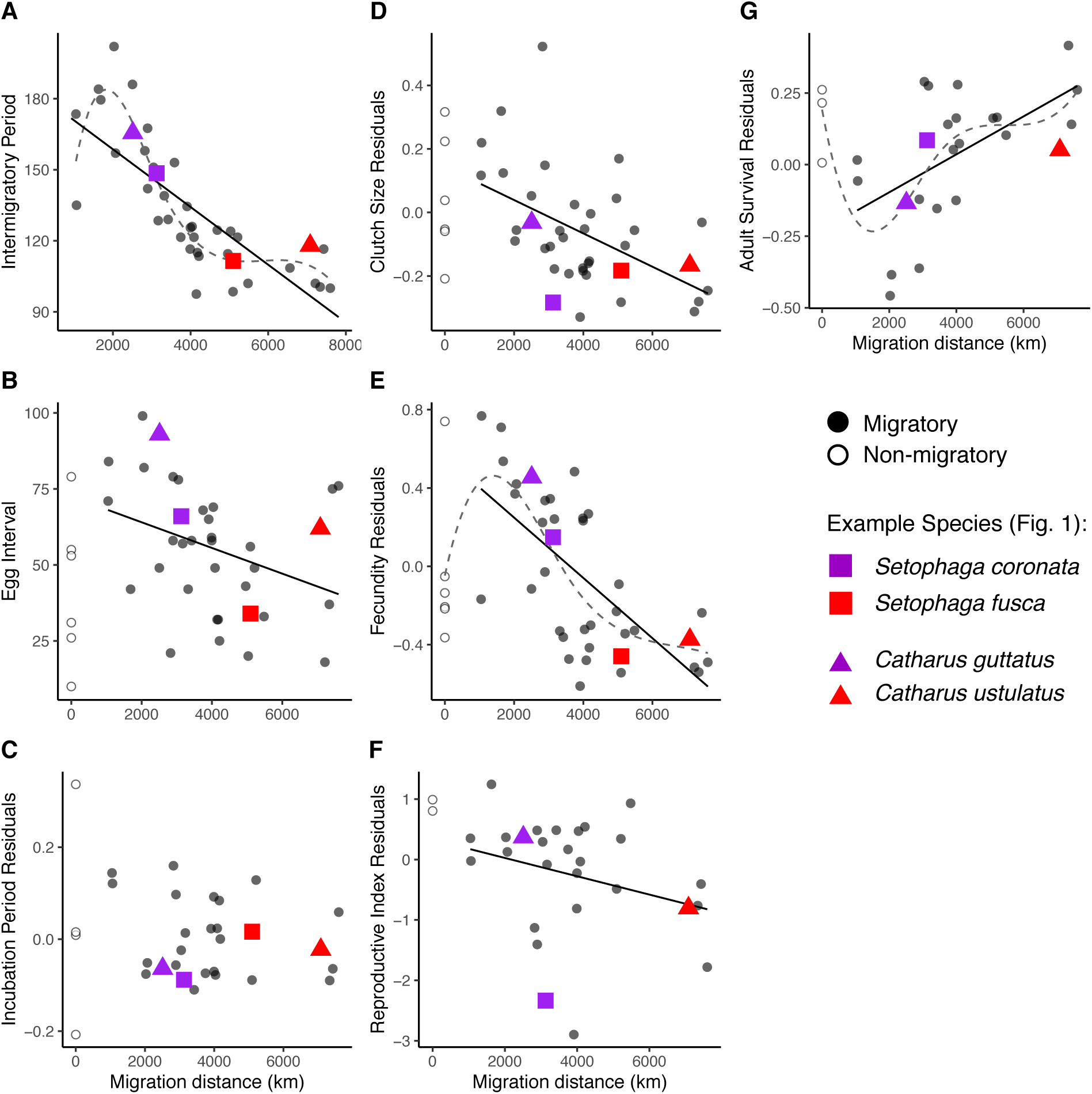
Relationships between migration distance and the variables (Table 1) related to breeding time allocation (A, B), developmental duration (C), reproductive output (D, E, F) and survival (F). For visualization purpose, the points shown in (C–G) are the residuals from a phylogenetic size-correction using the phyl.resid function in *phytools* (Revell 2009, 2012) because these panels represent variables thought to scale with body size. The solid line illustrates the linear relationship with migration distance among the 39 migratory species for variables in which migration distance was included in the best fit model (Table 2; all variables except incubation period, C). We include a cubic spline (dashed line) for plots with obvious non-linear patterns. For fecundity (E) and survival (G), this spline highlights that non-migratory species are more similar to long-distance migrants than to short-distance migrants, but that a relatively linear trend describes the relationship with these variables and migration distance across the migratory species. The splines also reveal that the steepest decline in annual fecundity (E) with migration distance mirror the steepest decline in intermigratory period (A), both occurring between migration distances of approximately 2000-4000km. Colored squares and triangles refer to the example species illustrated in Fig. 1, when data were available for each outcome variable. In (G) we did not plot the value for Golden-crowned Kinglet (*Regulus satrapa*), a short-distance migrant with a very low outlier value for survival, to more easily visualize the relationship among the remaining species.

This shortening of the breeding season does not seem to be associated with faster embryonic developmental rates for longer-distance migrants, as incubation period was not strongly correlated with migratory distance (Fig. 2) and the best model for incubation period contained only mass as a predictor (Tables 2, S2).

Average clutch size and annual fecundity were both significantly negatively associated with migratory distance (Fig. 2, Table 2). Reproductive Index (Desante et al. 2015) further suggested that the number of young fledged per adult trends negatively (but not significantly so) with increasing migratory distance across species (Fig. 2, Table 2). Apparent annual adult survival index (*ϕ*; Desante et al. 2015) was significantly positively associated with migratory distance (Table 2, Fig. 2).

### Migratory status and life history

In contrast to the relationships between migration distance and life history variation among the 39 migratory species, in which longer distance migrations were associated with “slower” life history values, non-migratory behavior (which involved 6 species that are year-round residents in a harsh winter environment with scarce resources) was associated with a shorter egg interval, lower clutch size and fecundity, and higher annual survival (Table 2, Fig. 2) than migratory species overall. That is, non-migratory species were more similar in survival and reproduction to long-distance migrants than to short-distance migrants, resulting in non-linear relationships with migration distance and some variables (Fig. 2).

### Winter climate and niche tracking

Migratory distance was significantly associated with winter climate PC1, indicating that species that migrate longer distances from the boreal region spend the winter in locations that are warmer, wetter and greener (Table 3, Fig. 3). However, migratory distance was also strongly negatively related to niche overlap (Zurell et al. 2018), indicating that longer-distance migrants have winter climates more dissimilar to their summer environments than do short-distance migrants (Table 3, Fig. 3).

**Table 3:** Results of climate and niche overlap models. Results of models testing how migration distance predicts winter climate (left panel) and how winter climate predicts adult survival (right panel). Coefficients with p < 0.05 are italicized and bolded. Models on the left use OLS whereas those on the right, which had significant values of λ (see Methods), use PGLS. Column “n” shows the number of species used in each model, based on data availability (Table S1). Winter climate is represented by a principle component value ranging from −2 (warmer, wetter, greener) to 2 (cooler, drier, less green). Niche overlap estimates comes from (Zurell et al. 2018), using their estimate that includes climate and NDVI and ranges from 0 to 0.42 in our dataset. Longer distance migrations are associated with wintering in warmer, wetter, and greener locations that show little niche overlap with the boreal breeding range. These conditions trend similarly with survival, but their effects are not significant in models with mass (Fig. 3).

| Response | Migration Distance | $\lambda$ | n | Response | Mass | Winter Climate PC1 | Niche Overlap (D) | $\lambda$ | n |
| --- | --- | --- | --- | --- | --- | --- | --- | --- | --- |
| Winter Climate PC1 | **-1.40** | 0 | 43 | Annual Adult Survival ($\phi$) | **0.44** | -0.08 | | 1.04 | 26 |
| Niche Overlap (D) | **-0.11** | 0 | 37 | Annual Adult Survival ($\phi$) | **0.51** | | -0.48 | 1.05 | 22 |

**Figure 3.**
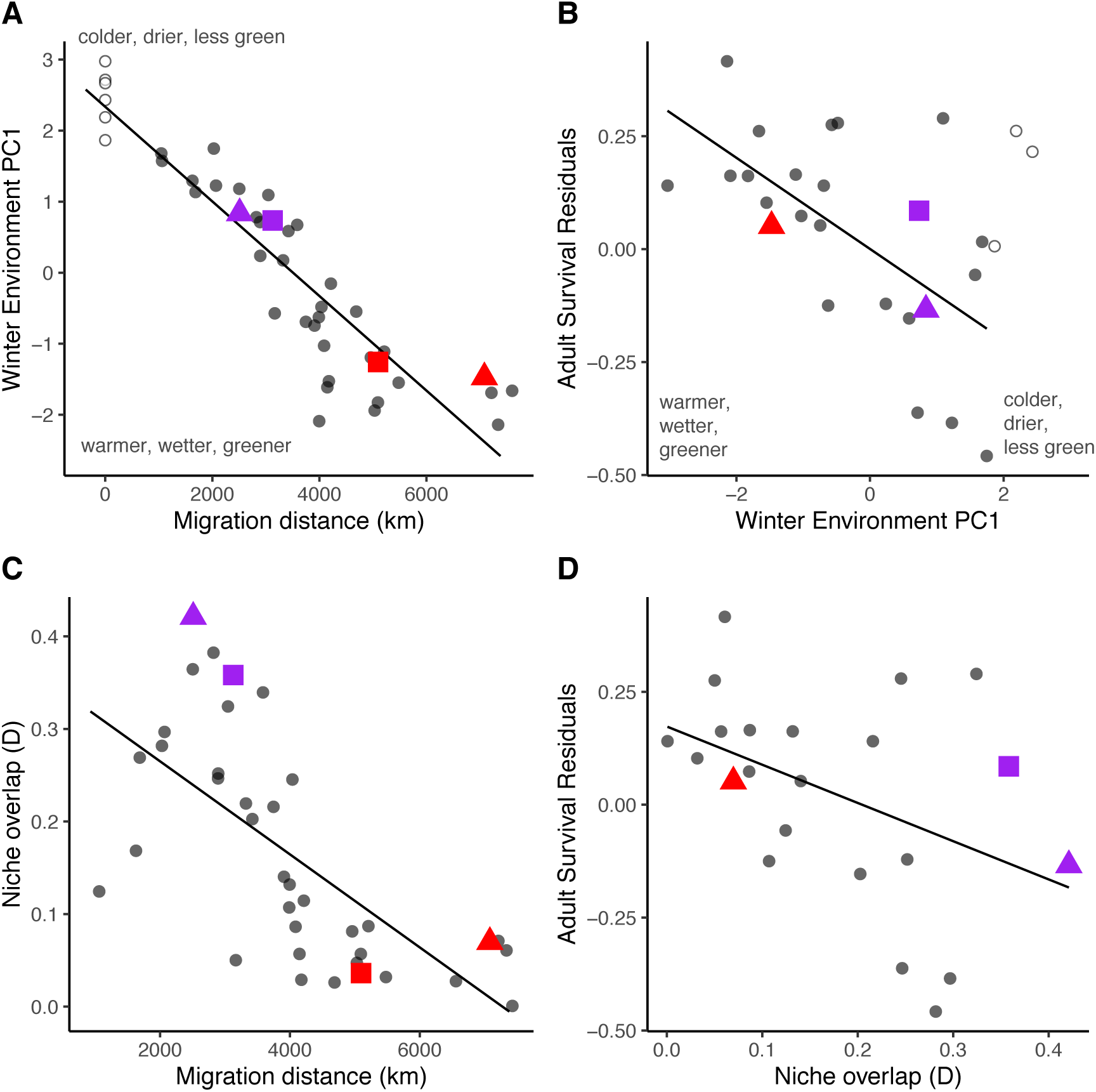
(A) Longer migrations carry species to winter ranges that are warmer, wetter and greener but (B) have less niche overlap (Zurell et al. 2018) with summer conditions. (C) Adult survival tends to be higher in warmer, wetter and greener winter ranges associated with long migrations (Fig. 2E) but that (D) have less overlap with breeding niches. Survival values are residuals from a phylogenetic size-correction (see Fig. 2). In (B, D) we did not plot the value for Golden-crowned Kinglet (*Regulus satrapa*), a short-distance migrant with a very low outlier value for survival, to more easily visualize the relationship among the remaining species. Niche overlap (D) were not calculated for the non-migratory species (Zurell et al. 2018). See Fig. 2 for color legend.

Although adult survival (*ϕ*) was significantly predicted by migratory distance, and migratory distance significantly predicted both winter climate PC1 and niche overlap, *ϕ* was not significantly predicted by either winter climate PC1 or niche overlap (Fig. 3, Table 3). Nevertheless, the trend of the relationship between *ϕ* and winter climate PC1 suggests that wintering in the warmer, wetter and greener environments associated with longer migrations may improve survival (Fig. 3, Table 3). Tracking a climatic niche throughout the year, however, did not improve survival.

## Discussion

Although there has been widespread interest in the consequences of life table parameters for the population demographics of migratory species (Sherry and Holmes 1995; Ricklefs 1997; Clark and Martin 2007; Faaborg et al. 2010), relatively little attention has been paid to the tradeoffs among these parameters across species with different migratory strategies and the evolutionary forces that shape them. Our focused analysis of demographic parameters from the boreal region shows that across 45 species of small-bodied, co-distributed birds breeding in similar habitats, seasonal migration has important and under-explored consequences for the trade-off between annual fecundity and survival. Further, these dynamics help explain the evolutionary dynamics underlying seasonal migration.

In support of the time allocation hypothesis (Greenberg 1980), our results indicate that migratory distance is negatively associated with temporal investment in breeding. As expected, migratory species that winter closer to their boreal breeding grounds migrate earlier in the spring and later in the fall and can be found incubating eggs during a longer time window than longer-distance migrants that winter closer to the equator, indicating a temporal constraint on breeding associated with longer-distance migration (Fig. 2). Long-distance migrants do not appear to compensate for this shorter breeding window through faster embryonic developmental times, which showed little relationship with migratory distance (Table 2, S2).

Our results further show that longer-distance migratory species have reduced clutch size and annual fecundity, as well as fewer fledged juveniles per adult (Desante et al. 2015), relative to shorter-distance migrants (Fig. 2). The relationship between annual fecundity and migration distance is consistent with, but stronger than, the relationship between clutch size and migration, because long-distance migrants in our study are not known to raise more than one successful brood in a single year (Table S1, Fig. 2). In some short-distance migratory species in our study the incidence of double brooding is likely low, but comparable data on incidence of multiple broods was lacking. Thus, future field studies are needed to determine whether the relationship between migration distance and reproductive output is closer to the curve for clutch size versus the steeper curve shown when maximum annual brood number is considered (Fig. 2).

Theory predicts that a decrease in annual fecundity across species should be associated with an increase in survival. By looking to the products of a long-term mark-recapture study with data specific to the boreal region (Desante et al. 2015), we found that migration distance trends positively with apparent annual adult survivorship in boreal-breeding birds. This is particularly striking given that the longest-distance migrants in our system undergo migrations of nearly 8000 km to equatorial forests (Fig. 1, Table S1) and migration is thought to be the costliest period of the annual cycle for survival in Neotropical migratory songbirds (Sillett and Holmes 2002). Yet, our results indicate that these long migrations provide a survival benefit compared to migrating shorter distances to higher latitude temperate and subtropical wintering latitudes (Fig. 2).

We further found that migratory distance and, to a lesser extent, annual adult survival were positively correlated with wintering in warmer, wetter and greener environments (Fig. 3). By contrast, the degree to which winter climate was similar to breeding climate (niche overlap) was negatively correlated with migratory distance. These results suggest that among the migratory species in our study, maximum survival is afforded to species that migrate the longest distances to humid equatorial forests, despite the energetic costs of long-distance migration and the fact that these long migrations lead to relatively greater differences in breeding season versus winter season climatic conditions. The importance of winter habitat for the annual cycle of migratory birds has been well documented within species, such as those whose individuals winter in habitats of differing quality (Norris et al. 2004; Faaborg et al. 2010; Rushing et al. 2016). However, the connection between migration distance, winter climate and annual survival, and its key evolutionary tradeoff with fecundity, has not been well elucidated in a macroecological comparative context. The specific macroecological conditions correlated with migration distance and winter survival in our study species, which are mainly forest-dwelling landbirds, are not likely to be generally applicable across migratory taxa with other habitat preferences and resource bases, but nevertheless these results speak broadly to the importance of winter resources for survival and reproduction.

Whereas long-distance migration is frequently described as costly to survival, our results show that the true cost of the longest migrations among migratory species in a temperate breeding community is a reduction in annual reproductive output. Temperate-breeding songbirds that endure migration of thousands of kilometers each year occupy a slower position on the slow-fast continuum of life history than their close relatives who migrate shorter distances but attempt to survive the winter closer to their breeding grounds. Species with shorter migrations to temperate or subtropical regions operate on the faster end of the slow-fast continuum, with relatively higher annual fecundity and lower adult survival rates. Aspects of this dynamic have been suggested previously in separate studies of survival or reproduction (Böhning-Gaese et al. 2000; Møller 2007; Bruderer and Salewski 2009; Conklin et al. 2017). However, a focused phylogenetic comparative analysis capable of revealing the evolutionary tradeoffs among these highly integrated life history variables has been lacking and thus the true relationship between long-distance migration, survival and fecundity has not been widely appreciated.

Our study also reveals meaningful distinctions between non-migratory species and short-distance migrants. We found that non-migratory boreal species had levels of reproductive output and survival more similar to long-distance migrants than to short-distance migrants (Fig. 2). Except for Canada Jay (*Perisoreus canadensis)*, all non-migratory species in our study (Table S1) are cavity-nesting chickadees (Paridae) or woodpeckers (Picidae), and all six species have specialized adaptations for surviving harsh boreal winters such as food-caching (Sherry 1989; Waite and Reeve 1993), social cooperation, or the ability to excavate grubs from trees. Our results therefore suggest a compelling pattern wherein the most specialized, extreme adaptations—required either to survive the harshest winters *in situ* or perform the longest migrations—exert strong selection pressures (Sol et al. 2010, 2016) that optimize annual adult survival at the cost of annual fecundity.

These results are consistent with emerging knowledge from field studies of some of the most extreme long-distance migratory birds, “long-jump” scolopacid sandpipers and godwits that breed in the high arctic and perform extreme flights to reach their wintering grounds (Conklin et al. 2017). In these species, there is increasing evidence that adult annual survival and longevity is surprisingly high given their remarkable annual journeys (Leyrer et al. 2013; Senner et al. 2019; Swift et al. 2020). This suggests that extreme long-distance migration selects strongly— potentially during the first year of life—for high quality individuals who as adults are capable of repeating these extraordinary feats every year, thus narrowing the “individual quality spectrum” to the highest quality individuals (Conklin et al. 2017). Our results provide an evolutionary life history framework for interpreting the high annual survival of these extreme long-distance migrants. Given the reproductive cost of long-distance migration, extreme migration can only be a successful evolutionary strategy if individual adult survival is high, thus placing these species on the slow end of the slow-fast continuum.

This perspective provides further evolutionary context for some surprising behaviors of migratory species. For example, one of the species in our study, the Veery (*Catharus fuscescens*), is a long-distance migrant breeding in temperate forests and wintering in Amazonia. A recent study (Heckscher 2018) found that Veeries shorten their breeding season in years with hurricane activity along their migratory route, putatively due to their perception of distant, future weather conditions. In other words, the evolution of their migration pattern—and the weather-detecting sensory systems that enable it—are so finely tuned to survival probability that some adult individuals will forego reproductive attempts to ensure a successful migration (Heckscher 2018). Our study shows that this species’ adaptive capacity to respond to subtle environmental conditions that threaten future survival during migration is representative of a point along a life history continuum wherein longer, more difficult migrations maximize annual survival at the cost of annual reproduction.

### Life history tradeoffs and the evolution of migration

The evolution of seasonal migration in birds has often been explained as an out-of-the-tropics process wherein species improve reproductive success by escaping the competition in the crowded tropics (Cox 1968, 1985; Rappole and Jones 2002). Support for the out-of-the tropics models has come in part from studies showing higher fecundity in temperate migrants versus tropical residents, which has been interpreted to mean that the evolution of migration facilitates greater reproductive success (reviewed in Salewski and Bruderer 2007; Winger et al. 2019*a*). A signature of this pattern is visible in some global studies where migration trends positively with fecundity variables (Jetz et al. 2008; Sibly et al. 2012). Although the tropical origins of long-distance migration have been contested in some lineages (Winger et al. 2014), other migratory lineages are ultimately of tropical biogeographic origin (Bruderer and Salewski 2008; Winger et al. 2019*a*). We have argued that even in lineages of tropical origin, migration does not evolve out of the tropics as a consequence of long-distance dispersal to improve reproductive success (Winger et al. 2019*a*). Rather, migration evolves when species expanding their ranges through normal dispersal processes encounter higher seasonality in a breeding location, often at a higher latitude, or when breeding populations face increases in seasonality through time *in situ* (Salewski and Bruderer 2007). The adaptive benefits of philopatry (returning to breed in similar locations as one was reared; Davis and Stamps 2004) select for individuals that return to their breeding grounds in spring (Winger et al. 2019*a*). From this perspective, migration is an adaptive survival strategy that allows for persistence in seasonal areas, regardless of the biogeographic origin of a lineage.

Our study lends support to this survival hypothesis by demonstrating the relationship between migration, reproduction and survival among a community of species that experiences high seasonality in the breeding range. Overall, the annual fecundity of the migratory species in our study is likely greater than that of close tropical-breeding relatives, since it varies along a strong latitudinal gradient in clutch size (Ricklefs 2000*a*; Clark and Martin 2007). However, annual fecundity is also relatively high among the non-migratory species that occupy temperate latitudes. Thus, the evolution of migration promotes fecundity only insofar as it promotes survival and persistence of populations in highly seasonal environmental conditions where annual fecundity—and mortality—are generally higher than in the tropics. In other words, the colonization of temperate, seasonal environments moves populations towards a “faster” life history compared to tropical species (Martin 2015), where high fecundity trades off with high mortality. The evolution of long-distance migration mitigates this high annual mortality by bolstering winter survival, but at a necessary cost to annual fecundity, thereby ‘slowing down’ life history. The longer the migration, the slower the life history.

We suggest that the evolution of seasonal migration should be regarded as a fundamental life history tradeoff intimately connected to environmentally driven patterns of survival and fecundity, as opposed to a unique strategy with a deterministic benefit for reproductive output.

### Opportunities for further research

Our analyses of data compiled from published sources highlight the potential for further insights to be gained from focused field studies. First, reliable data on longevity across our study species is important to understand lifetime reproductive success as opposed to annual patterns. Second, although annual survival is thought to be age-independent in adult birds (Ricklefs 1997), age-specific life tables of survival and fecundity will improve our understanding of how migratory behavior influences the slow-fast continuum and the consequences for population demography (Saether and Bakke 2000). Data on juvenile survival rates, while much more difficult to study due to the likelihood of conflating mortality with natal dispersal, will help illuminate the differences in selection pressures facing species across the migratory spectrum (Conklin et al. 2017)

An important mystery highlighted by our study concerns the post-breeding, pre-migratory periods, which is in general the most poorly understood period of the avian annual cycle (Clark and Martin 2007). Even if post-embryonic developmental rates are longer in shorter-distance migrants, short-distance migrants still spend substantially greater time post-development on or near their breeding sites before fall migration than do long-distance migrants (Fig. 4). That is, short-distance migrants have a “faster” life history despite an annual schedule that is less temporally constrained. What do hatch-year birds in short-distance migratory and resident species—which will not breed until the following spring—do with their “extra” time (Fig. 4)?

**Figure 4.**
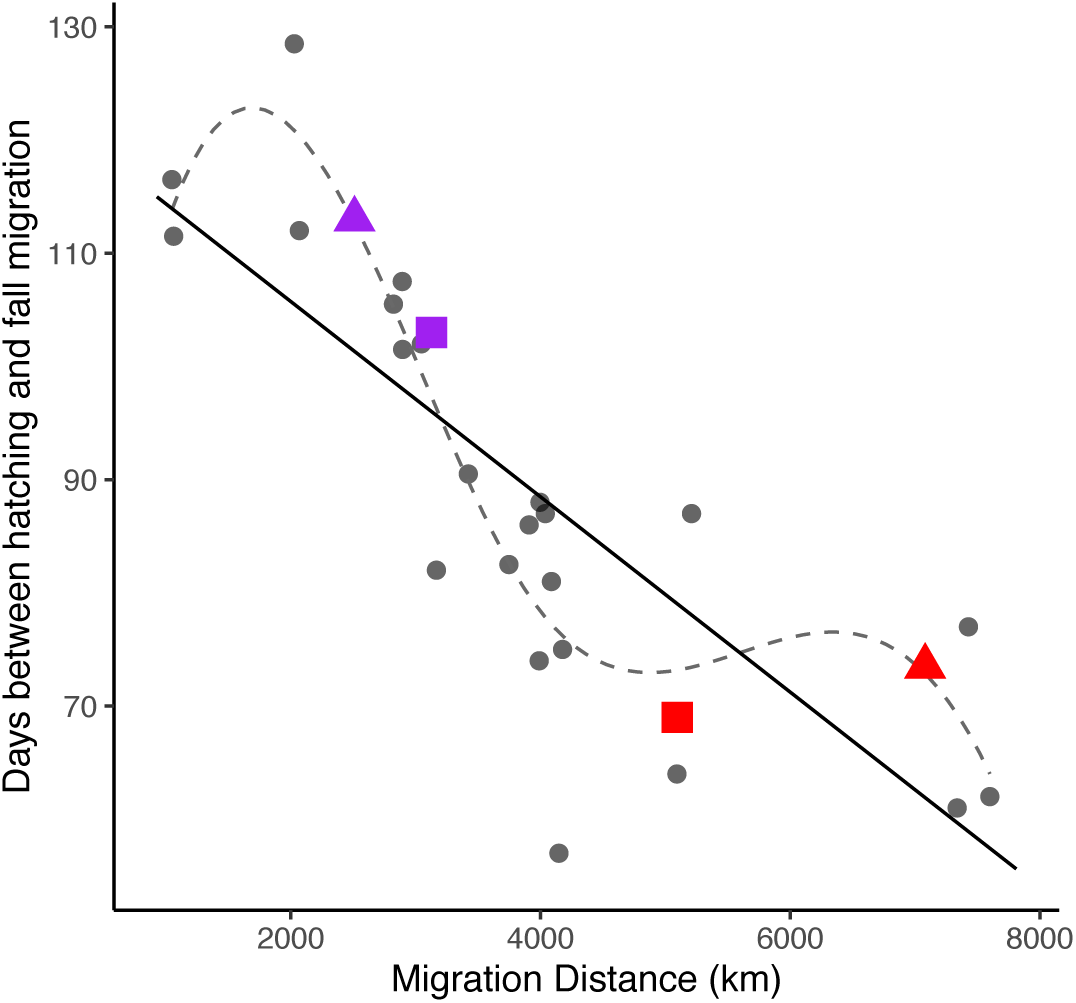
Duration of time between hatching and fall migration across the spectrum of migratory distance. Hatching date was estimated as the median of early dates that nests have been found with eggs in Ontario for each species from *Nidiology* (Peck and James 1983, 1987) plus average incubation period for 27 species with data on incubation period from *Nidiology*. Fall migration was estimated from eBird as the median date when each species passes through Chicago, IL, a stopover site close to the southern edge of the boreal region (see Methods), meant to approximate the beginning of fall migration. Short-distance migrants spend substantially greater time on or near their breeding grounds between hatching and fall migration than long-distance migrants. Data were not available on the durations of the nestling and fledgling stages (post-embryonic development) of each species, but differences in post-embryonic developmental rates are unlikely to fill this extra time. Incubation period was not correlated with migration distance (Fig. 2), and thus the relationship between migration distance and days between clutch initiation and fall migration is nearly identical (not shown). After the fledgling period, birds are thought to undergo prospecting for future breeding sites, an important phase of natal dispersal, which may allow them to settle on their first breeding territory more quickly following spring migration. We propose that hatch year short-distance migrants may use their greater time on the breeding grounds for more extensive prospecting.

One possibility is that shorter-distance migratory species and residents invest more time in the prospecting phase of dispersal (Reed et al. 1999), wherein they use the pre-migratory period to search nearby breeding locations for future breeding sites (Cormier and Taylor 2019). By contrast, long-distance migratory species, with constrained time for prospecting, may be under greater selection to return to familiar breeding locations (Winger et al. 2019*a*), which could counterintuitively constrain dispersal distances in longer-distance migrants. This possibility is important to investigate because annual survival estimates based on mark-recapture could be unreliable if adult dispersal rates differ systematically across species.

### Implications for the conservation of migratory birds

Our study shows that at its core, migration is a survival strategy that evolves when the benefits of escaping the breeding region during the resource-poor season outweigh the risks of long-distance journeys. Yet, this strategy is only likely to be beneficial when suitable habitat is available on the wintering grounds and throughout migration (Conklin et al. 2017). It is obvious that the pace at which anthropogenic habitat destruction has occurred on the Neotropical wintering grounds and at stopover sites for the long-distance migrants in our study far exceeds the ability of these species to adjust their delicately balanced life history strategies. Other major disruptions to migration, such as from artificial light (Loss et al. 2015; Winger et al. 2019*b*), may further tip the scales against long-distance migration as an effective strategy. Biologists have warned for decades that long-distance migratory species are among the species most at risk of population decline in the Anthropocene (Robbins et al. 1989; Rosenberg et al. 2019). Our study, by highlighting that lowered fecundity is a cost of the heightened survival afforded to long-distance migrants, provides an evolutionary context for understanding the importance of habitat conservation and safe passage for migratory birds throughout their journeys.

## Acknowledgments

We thank James Saracco and David DeSante for helpful advice regarding *Vital Rates* parameters and for making these data available. We thank Jacob Berv, Eric Gulson-Castillo, Theunis Piersma, Kristen Wacker and Marketa Zimova for helpful feedback on earlier versions of the manuscript.

## Figures and Tables

**Table S1:**
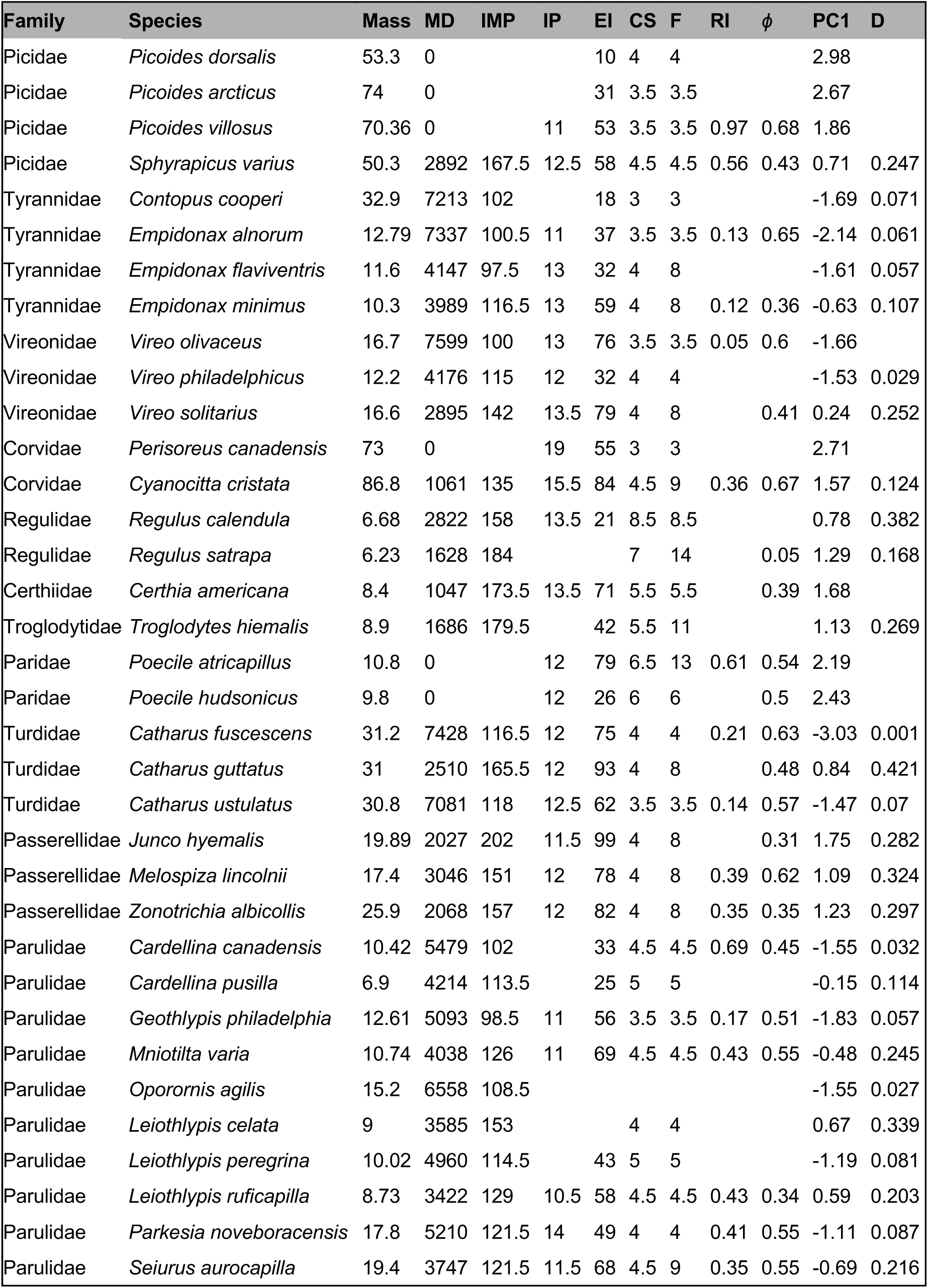
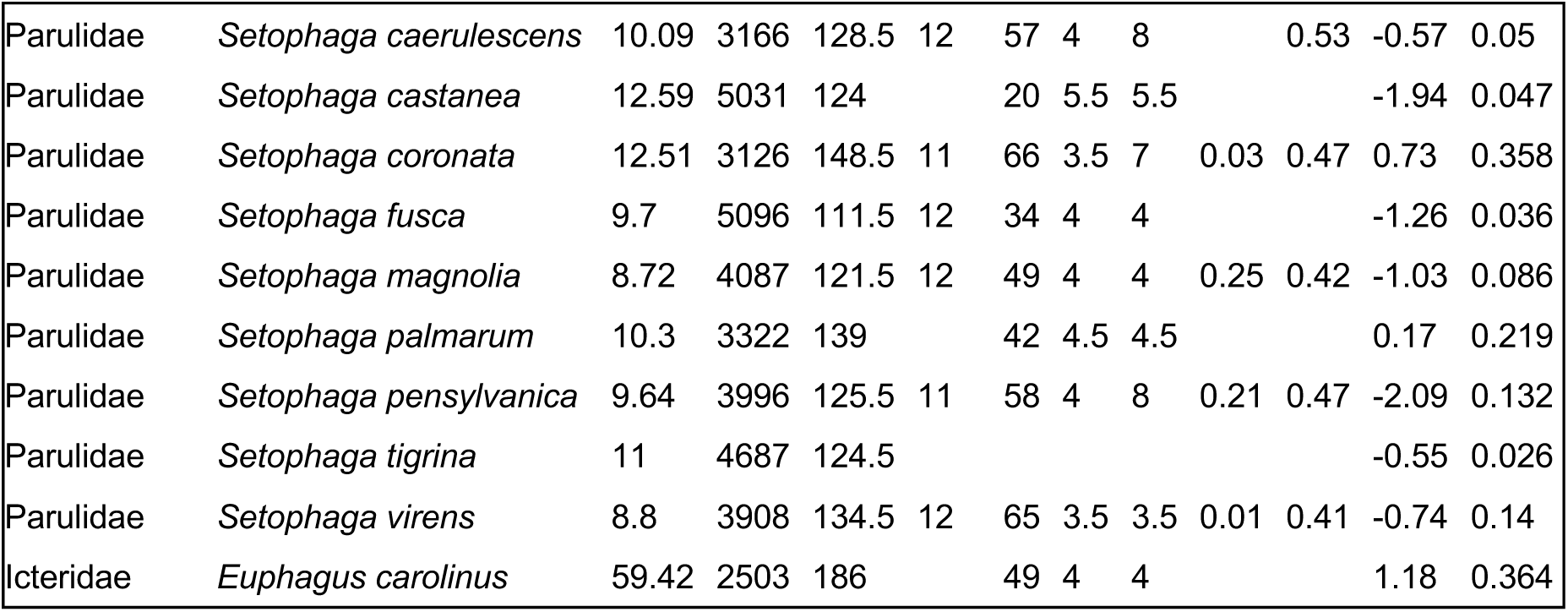
Species and trait data used in analyses. Species and data used in analyses. MD: Migration distance (km); IMP: intermigratory period (days); IP: incubation period (days); EI: egg interval (days); CS: average clutch size; F: fecundity; RI: Reproductive Index; *ϕ:* Annual Adult Survival; PC1: first principal component of PCA of temperature, precipitation and NDVI on winter ranges; D: niche overlap value (Zurell et al. 2018). See Table 1 and Methods for descriptions and data sources for variables.

**Table S2:**
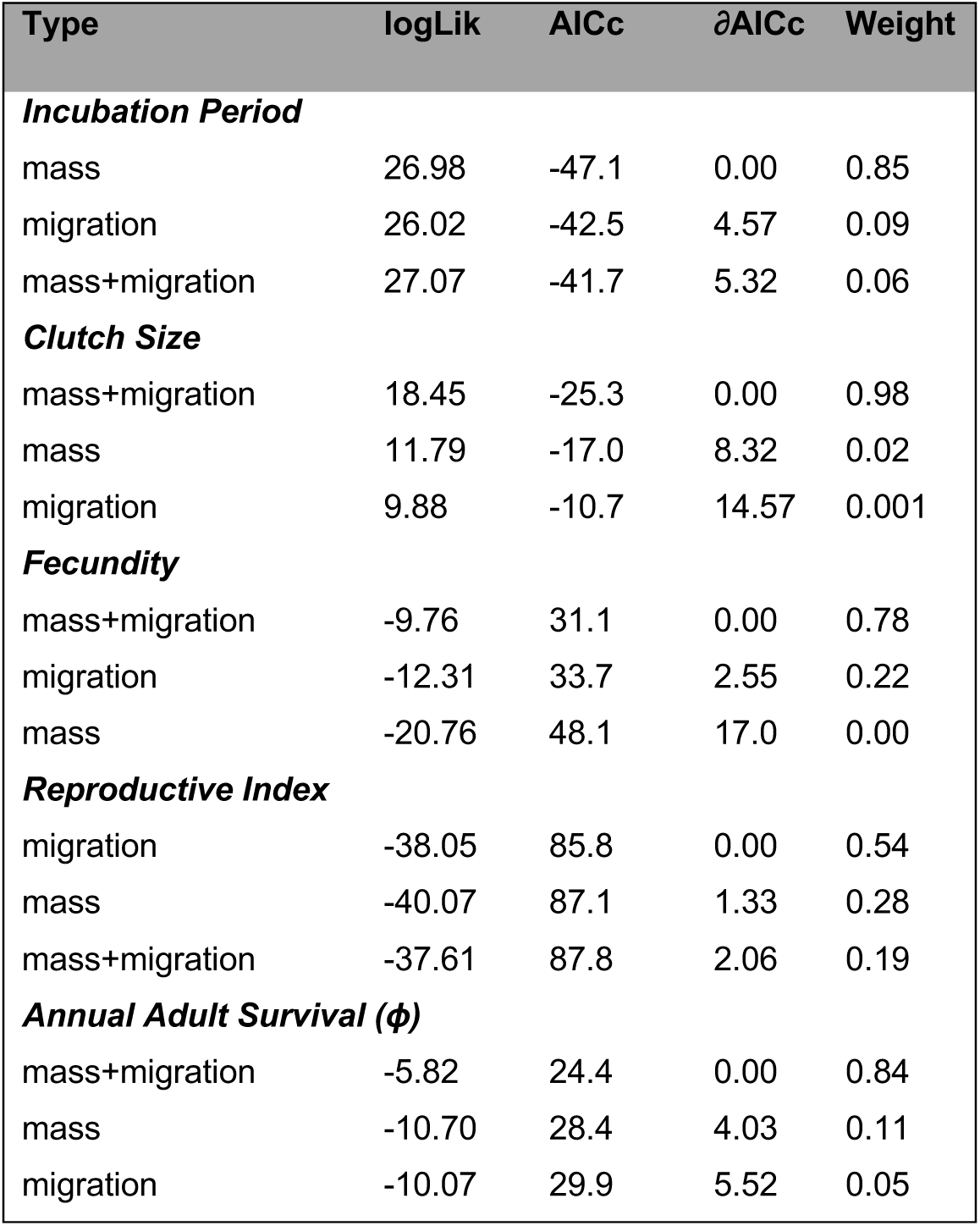
Results of AICc model comparison. Results of model selection for each life history variable in which models with mass and migration variables were compared. For all response variables except incubation period, models with migration predictors (status and distance) outperform those without. Model selection was performed on models using either OLS or PGLS depending on the phylogenetic significance of the residuals of a full model (Table 2). Outcome variables and mass were log-transformed.

| Type | logLik | AICc | ΔAICc | Weight |
| --- | --- | --- | --- | --- |
| ***Incubation Period*** |  |  |  |  |
| mass | 26.98 | -47.1 | 0.00 | 0.85 |
| migration | 26.02 | -42.5 | 4.57 | 0.09 |
| mass+migration | 27.07 | -41.7 | 5.32 | 0.06 |
| ***Clutch Size*** |  |  |  |  |
| mass+migration | 18.45 | -25.3 | 0.00 | 0.98 |
| mass | 11.79 | -17.0 | 8.32 | 0.02 |
| migration | 9.88 | -10.7 | 14.57 | 0.001 |
| ***Fecundity*** |  |  |  |  |
| mass+migration | -9.76 | 31.1 | 0.00 | 0.78 |
| migration | -12.31 | 33.7 | 2.55 | 0.22 |
| mass | -20.76 | 48.1 | 17.0 | 0.00 |
| ***Reproductive Index*** |  |  |  |  |
| migration | -38.05 | 85.8 | 0.00 | 0.54 |
| mass | -40.07 | 87.1 | 1.33 | 0.28 |
| mass+migration | -37.61 | 87.8 | 2.06 | 0.19 |
| ***Annual Adult Survival ($\phi$)*** |  |  |  |  |
| mass+migration | -5.82 | 24.4 | 0.00 | 0.84 |
| mass | -10.70 | 28.4 | 4.03 | 0.11 |
| migration | -10.07 | 29.9 | 5.52 | 0.05 |

**Table S3:**
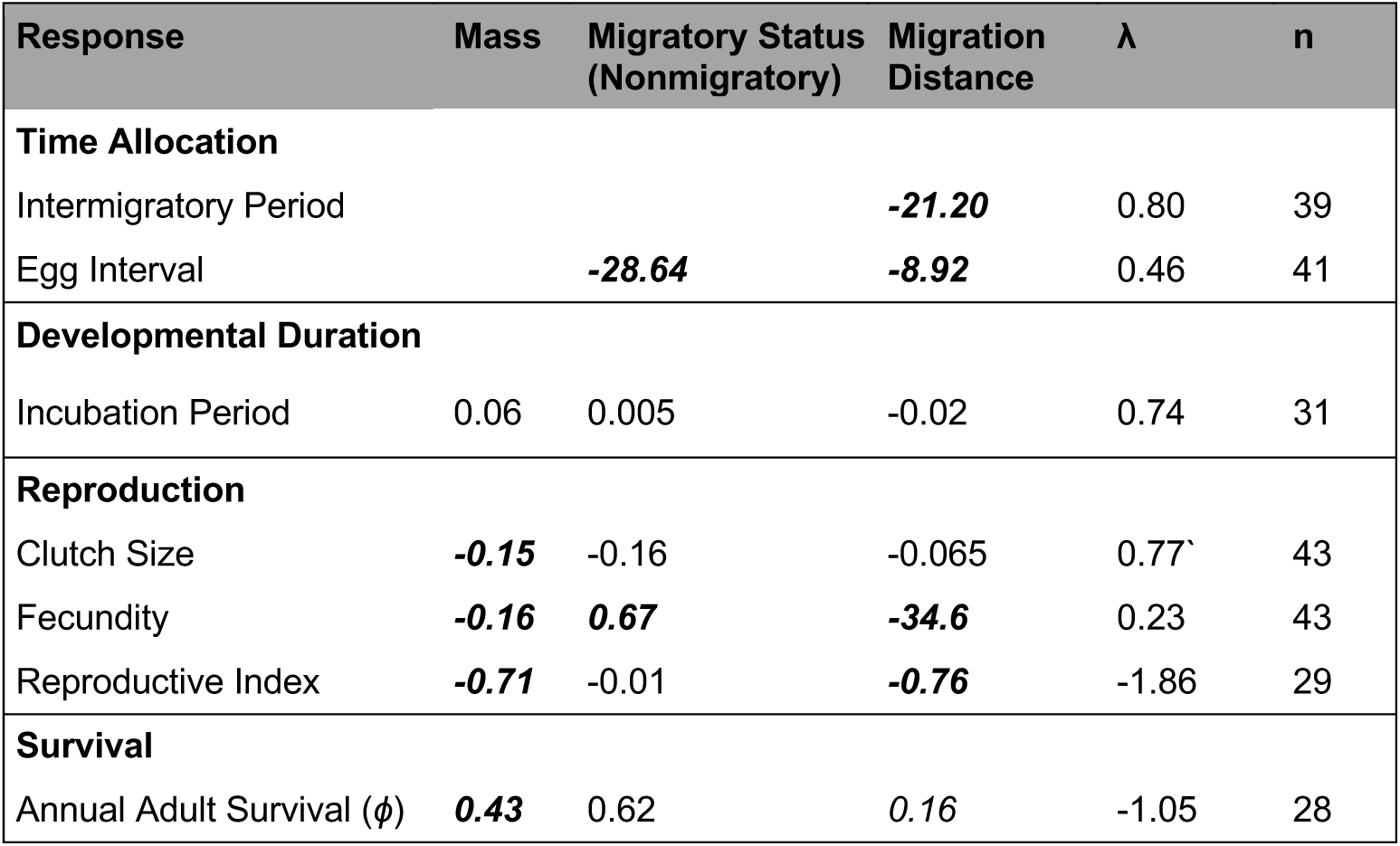
Results of PGLSλ models. Summary of full PGLS models for each life history variable. For each response, we fit a PGLS model which optimizes an estimate of λ simultaneously with model fitting (Revell 2010). The trends are consistent with those reported in Table 2 using OLS, though not all trends are significant in PGLS. Coefficients with p < 0.05 are italicized and bolded.

| Response | Mass | Migratory Status<br>(Nonmigratory) | Migration<br>Distance | $\lambda$ | n |
| --- | --- | --- | --- | --- | --- |
| **Time Allocation** |  |  |  |  |  |
| Intermigratory Period |  |  | **-21.20** | 0.80 | 39 |
| Egg Interval |  | **-28.64** | **-8.92** | 0.46 | 41 |
| **Developmental Duration** |  |  |  |  |  |
| Incubation Period | 0.06 | 0.005 | -0.02 | 0.74 | 31 |
| **Reproduction** |  |  |  |  |  |
| Clutch Size | **-0.15** | -0.16 | -0.065 | 0.77 | 43 |
| Fecundity | **-0.16** | **0.67** | **-34.6** | 0.23 | 43 |
| Reproductive Index | **-0.71** | -0.01 | **-0.76** | -1.86 | 29 |
| **Survival** |  |  |  |  |  |
| Annual Adult Survival ($\phi$) | **0.43** | 0.62 | 0.16 | -1.05 | 28 |

## References

Alves, J. A., T. G. Gunnarsson, D. B. Hayhow, G. F. Appleton, P. M. Potts, W. J. Sutherland, and J. A. Gill. 2013. Costs, benefits, and fitness consequences of different migratory strategies. Ecology 94:11–17.

Bartón, K. 2019. MuMIn: Multi-Model Inference.

Billerman, S., B. Keeney, P. Rodewald, and T. Schulenberg, eds. 2020. Birds of the World. Cornell Laboratory of Ornithology, Ithaca, NY.

Bird, J. P., R. Martin, H. R. Akçakaya, J. Gilroy, I. J. Burfield, S. Garnett, A. Symes, et al. 2020. Generation lengths of the world’s birds and their implications for extinction risk. Conservation Biology In Press.

Bivand, R., T. Keitt, and B. Rowlingson. 2019. rgdal: Bindings for the “Geospatial” Data Abstraction Library.

Bivand, R., and C. Rundel. 2019. rgeos: Interface to Geometry Engine - Open Source (‘GEOS’).

Böhning-Gaese, K., B. Halbe, N. Lemoine, and R. Oberrath. 2000. Factors influencing the clutch size, number of broods and annual fecundity of North American and European land birds. Evolutionary Ecology Research 2:823–839.

Bruderer, B., and V. Salewski. 2008. Evolution of bird migration in a biogeographical context. Journal of Biogeography 35:1951–1959.

Bruderer, B. 2009. Lower annual fecundity in long-distance migrants than in less migratory birds of temperate Europe. Journal of Ornithology 150:281–286.

Buchan, C., J. J. Gilroy, I. Catry, and A. M. A. Franco. 2019. Fitness consequences of different migratory strategies in partially migratory populations: A multi-taxa meta-analysis. Journal of Animal Ecology 89:678–690.

Bird Studies Canada and NABCI. 2014. Bird Conservation Regions. Published by Bird Studies Canada on behalf of the North American Bird Conservation Initiative.

Clark, M. E., and T. E. Martin. 2007. Modeling tradeoffs in avian life history traits and consequences for population growth. Ecological Modelling 209:110–120.

Conklin, J. R., N. R. Senner, P. F. Battley, and T. Piersma. 2017. Extreme migration and the individual quality spectrum. Journal of Avian Biology 48:19–36.

Cooney, C. R., C. Sheard, A. D. Clark, S. D. Healy, A. Liker, S. E. Street, C. A. Troisi, et al. 2020. Ecology and allometry predict the evolution of avian developmental durations. Nature Communications 11:2383.

Cormier, D. A., and P. D. Taylor. 2019. Contrasting patterns of post-fledging movements of two sympatric warbler species with different life-history strategies. Journal of Avian Biology 50:1–11.

Cox, G. 1985. The evolution of avian migration systems between the temperate and tropical regions of the New World. American Naturalist 126:451–474.

Cox, G. W. 1968. The role of competition in the evolution of migration. Evolution 22:180–192.

Davis, J. M., and J. A. Stamps. 2004. The effect of natal experience on habitat preferences. Trends in Ecology and Evolution 19:411–416.

Desante, D. F., D. R. Kaschube, and J. F. Saracco. 2015. Vital Rates of North American Landbirds. The Institute for Bird Populations.

Dunning, J. B. J. 2008. CRC Handbook of Avian Body Masses. CRC Press, Boca Raton.

Ely, C. R., and B. W. Meixell. 2015. Demographic outcomes of diverse migration strategies assessed in a metapopulation of tundra swans. Movement Ecology 4:1–15.

Faaborg, J., R. T. Holmes, A. D. Anders, K. L. Bildstein, K. M. Dugger, S. A. Gauthreaux Jr, P. Heglund, et al. 2010. Recent advances in understanding migration systems of New World land birds. Ecological Monographs 80:3–48.

Fick, S. E., and R. J. Hijmans. 2017. WorldClim 2: new 1–km spatial resolution climate surfaces for global land areas. International Journal of Climatology 37:4302–4315.

Gasser, M., M. Kaiser, D. Berrigan, and S. C. Stearns. 2000. Life-history correlates of evolution under high and low adult mortality. Evolution 54:1260–1272.

Gómez, C., E. A. Tenorio, P. Montoya, and C. D. Cadena. 2016. Niche-tracking migrants and niche-switching residents: evolution of climatic niches in New World warblers (Parulidae). Proceedings of the Royal Society B: Biological Sciences 283:20152458.

Greenberg, R. 1980. Demographic Aspects of Long-distance Migration. Pages 493–504 in A. Keast and E. Morton, eds. Migrant Birds in the Neotropics. Smithsonian Institution Press, Washington, D.C.

Greenberg, R., and P. P. Marra. 2005. Birds of Two Worlds: The Ecology and Evolution of Migration.

Heckscher, C. M. 2018. A Nearctic-Neotropical migratory songbird’s nesting phenology and clutch size are predictors of accumulated cyclone energy. Scientific Reports 8:1–6.

Hijmans, R. J. 2019. geosphere: Spherical Trigonometry. R package.

Hostetler, J. A., T. S. Sillett, and P. P. Marra. 2015. Full-annual-cycle population models for migratory birds. The Auk 132:433–449.

BirdLife International, and Naturserve. 2014. Bird species distribution maps of the world. Cambridge, UK.

Jetz, W., C. H. Sekercioglu, and K. Böhning-Gaese. 2008. The worldwide variation in avian clutch size across species and space. Plos One 6:2650–2657.

Jetz, W., G. H. Thomas, J. B. Joy, K. Hartmann, and A. O. Mooers. 2012. The global diversity of birds in space and time. Nature 491:444–448.

Karagicheva, J., E. Rakhimberdiev, A. Saveliev, and T. Piersma. 2018. Annual chronotypes functionally link life histories and life cycles in birds. Functional Ecology 32:2369–2379.

Kardynal, K. J., and K. A. Hobson. 2017. The pull of the Central Flyway? Veeries breeding in western Canada migrate using an ancestral eastern route. Journal of Field Ornithology 88:262–273.

Kelly, J., and R. Hutto. 2005. An east-west comparison of migration in North American wood warblers. Condor 107:197–211.

Ketterson, E. D., and V. Nolan. 1982. The role of migration and winter mortality in the life history of a temperate-zone migrant, the Dark-Eyed Junco, as determined from demographic analyses of winter populations. The Auk 99:243–259.

Leyrer, J., T. Lok, M. Brugge, B. Spaans, B. K. Sandercock, and T. Piersma. 2013. Mortality within the annual cycle: Seasonal survival patterns in Afro-Siberian Red Knots *Calidris canutus* canutus. Journal of Ornithology 154:933–943.

Lok, T., O. Overdijk, and T. Piersma. 2015. The cost of migration: Spoonbills suffer higher mortality during trans-Saharan spring migrations only. Biology Letters 11:20140944.

Lok, T., L. Veldhoen, O. Overdijk, J. M. Tinbergen, and T. Piersma. 2017. An age-dependent fitness cost of migration? Old trans-Saharan migrating spoonbills breed later than those staying in Europe, and late breeders have lower recruitment. Journal of Animal Ecology 86:998–1009.

Loss, S. R., T. Will, and P. P. Marra. 2015. Direct mortality of birds from anthropogenic causes. Annual Review of Ecology, Evolution, and Systematics 46:99–120.

Marra, P., E. B. Cohen, S. R. Loss, J. E. Rutter, and C. M. Tonra. 2015. A call for full annual cycle research in animal ecology. Biology Letters 11:2015.0552.

Martin, T. E. 2002. A new view of avian life-history evolution tested on an incubation paradox. Proceedings of the Royal Society B: Biological Sciences 269:309–316.

Martin, T. E. 2004. Avian life-history evolution has an eminent past: does it have a bright future? The Auk 121:289–301.

Martin, T. E. 2015. Age-related mortality explains life history strategies of tropical and temperate songbirds. Science 349:966–970.

Møller, A. P. 2007. Senescence in relation to latitude and migration in birds. Journal of Evolutionary Biology 20:750–757.

Mönkkönen, M. 1992. Life history traits of Palaearctic and Nearctic migrant passerines. Ornis Fennica 69:161–172.

Muñoz, A. P., M. Kéry, P. V. Martins, and G. Ferraz. 2018. Age effects on survival of Amazon forest birds and the latitudinal gradient in bird survival. The Auk 135:299–313.

Newton, I. 2007. Weather-related mass-mortality events in migrants. Ibis 149:453–467.

Norris, D. R., P. P. Marra, T. K. Kyser, T. W. Sherry, and L. M. Ratcliffe. 2004. Tropical winter habitat limits reproductive success on the temperate breeding grounds in a migratory bird. Proceedings of the Royal Society B: Biological Sciences 271:59–64.

Omernik, J. M., and G. E. Griffith. 2014. Ecoregions of the Conterminous United States: Evolution of a Hierarchical Spatial Framework. Environmental Management 54:1249–1266.

Paradis, E., and K. Schliep. 2019. Ape 5.0: An environment for modern phylogenetics and evolutionary analyses in R. Bioinformatics 35:526–528.

Peck, G. W., and R. D. James. 1983. Breeding Birds of Ontario: Nidiology and Distribution Volume 1 Nonpasserines. Royal Ontario Museum, Toronto, Canada.

Peck, G. W. 1987. Breeding Birds of Ontario: Nidiology and Distribution Volume 2 Passerines. Royal Ontario Museum, Toronto, Canada.

Pegan, T. M., and B. M. Winger. 2020. The influence of seasonal migration on range size in temperate North American passerines. Ecography In Press.

Pinheiro, J., D. Bates, S. DebRoy, D. Sarkar, and R Core Team. 2019. nlme: Linear and Nonlinear Mixed Effects Models. https://CRAN.R-project.org/package=nlme.

Pinzon, J. E., and C. J. Tucker. 2014. A non-stationary 1981-2012 AVHRR NDVI3g time series. Remote Sensing 6:6929–6960.

Rappole, J. H., and P. Jones. 2002. Evolution of old and new world migration systems. Ardea 90:525–537.

Reed, J. M., T. Boulinier, E. Danchin, and L. W. Oring. 1999. Informed dispersal: Prospecting by birds for breeding sites. Current Ornithology 15:189–259.

Revell, L. J. 2009. Size-correction and principal components for interspecific comparative studies. Evolution 63:3258–3268.

Revell, L. J. 2010. Phylogenetic signal and linear regression on species data. Methods in Ecology and Evolution 1:319–329.

Revell, L. J. 2012. phytools: An R package for phylogenetic comparative biology (and other things). Methods in Ecology and Evolution 3:217–223.

Reznick, D. 1984. Costs of reproduction?: An evaluation of the empirical evidence. Oikos 44:257–267.

Ricklefs, R. E. 1980. Comparative avian demography. Pages 1–32 in R. F. Johnston, ed. Current Ornithology Vol. 1. Plenum Press, Lawrence, KS.

Ricklefs, R. E. 1997. Comparative demography of New World populations of thrushes (Turdus spp.). Ecological Monographs 67:23–43.

Ricklefs, R. E. 2000a. Density dependence, evolutionary optimization, and the diversification of avian life histories. The Condor 102:9–22.

Ricklefs, R. E. 2000b. Lack, Skutch, and Moreau: The early development of life-history thinking. Condor 102:3–8.

Ricklefs, R. E., and M. Wikelski. 2002. The physiology/life-history nexus. TRENDS in Ecology & Evolution 17:462–468.

Robbins, C. S., J. R. Sauer, R. S. Greenberg, and S. Droege. 1989. Population declines in North American birds that migrate to the neotropics. Proceedings of the National Academy of Sciences USA 86:7658–7662.

Rosenberg, K. V, A. M. Dokter, P. J. Blancher, J. R. Sauer, A. C. Smith, P. A. Smith, J. C. Stanton, et al. 2019. Decline of the North American avifauna. Science 366:120–124.

Rushing, C. S., P. P. Marra, and M. R. Dudash. 2016. Winter habitat quality but not long-distance dispersal influences apparent reproductive success in a migratory bird. Ecology 97:1218–1227.

Saether, B.-E. 1988. Pattern of covariation between life-history traits of European birds. Nature 331:616–617.

Saether, B.-E., and O. Bakke. 2000. Avian life history variation and contribution of demographic traits to the population growth rate. Ecology 81:642–653.

Salewski, V., and B. Bruderer. 2007. The evolution of bird migration—a synthesis. Naturwissenschaften 94:268–279.

Senner, N. R., M. A. Verhoeven, J. M. Abad-Gómez, J. A. Alves, J. C. E. W. Hooijmeijer, R. A. Howison, R. Kentie, et al. 2019. High migratory survival and highly variable migratory behavior in black-tailed godwits. Frontiers in Ecology and Evolution 7:1–11.

Sherry, F. 1989. Food storing in the Paridae. Wilson Bulletin 101:289–304.

Sherry, T. W., and R. T. Holmes. 1995. Summer versus winter limitation of populations: what are the issues and what is the evidence? Pages 85–120 in T. E. Martin and D. M. Finch, eds. Ecology and Management of Neotropical Migratory Birds. Oxford University Press, New York.

Sibly, R. M., C. C. Witt, N. A. Wright, C. Venditti, W. Jetz, and J. H. Brown. 2012. Energetics, lifestyle, and reproduction in birds. Proceedings of the National Academy of Sciences of the United States of America 109:10937–10941.

Sillett, T. S., and R. T. Holmes. 2002. Variation in survivorship of a migratory songbird throughout its annual cycle. Journal of Animal Ecology 71:296–308.

Sol, D., N. Garcia, A. Iwaniuk, K. Davis, A. Meade, W. A. Boyle, and T. Székely. 2010. Evolutionary divergence in brain size between migratory and resident birds. PLoS ONE 5:1–8.

Sol, D., F. Sayol, S. Ducatez, and L. Lefebvre. 2016. The life-history basis of behavioural innovations. Philosophical Transactions of the Royal Society B: Biological Sciences 371:20150187.

Somveille, M., A. S. L. Rodrigues, and A. Manica. 2018. Energy efficiency drives the global seasonal distribution of birds. Nature Ecology and Evolution 2:962–969.

Somveille, M., A. Manica, and A. S. Rodrigues. 2019. Where the wild birds go: explaining the differences in migratory destinations across terrestrial bird species. Ecography 42:225–236.

Stearns, S. C. 1976. Life-history tactics: a review of the ideas. The Quarterly Review of Biology 51:3–47.

Sullivan, B. L., C. L. Wood, M. J. Iliff, R. E. Bonney, D. Fink, and S. Kelling. 2009. eBird: A citizen-based bird observation network in the biological sciences. Biological Conservation 142:2282–2292.

Swanson, D. L., and T. Garland. 2009. The evolution of high summit metabolism and cold tolerance in birds and its impact on present-day distributions. Evolution 63:184–194.

Swift, R. J., A. D. Rodewald, J. A. Johnson, B. A. Andres, and N. R. Senner. 2020. Seasonal survival and reversible state effects in a long-distance migratory shorebird. Journal of Animal Ecology In Press.

Townsend, A. K., T. S. Sillett, N. K. Lany, S. A. Kaiser, N. L. Rodenhouse, M. S. Webster, and R. T. Holmes. 2013. Warm springs, early lay dates, and double brooding in a North American migratory songbird, the Black-Throated Blue Warbler. PLoS ONE 8:e59467.

Waite, T. A., and J. D. Reeve. 1993. Food storage in Gray Jays: Source type and cache dispersion. Ethology 93:326–336.

Winger, B. M., F. K. Barker, and R. H. Ree. 2014. Temperate origins of long-distance seasonal migration in New World songbirds. Proceedings of the National Academy of Sciences of the United States of America 111:12115–12120.

Winger, B. M., G. G. Auteri, T. M. Pegan, and B. C. Weeks. 2019a. A long winter for the Red Queen: rethinking the evolution of seasonal migration. Biological Reviews 94:737–752.

Winger, B. M., B. C. Weeks, A. Farnsworth, A. W. Jones, M. Hennen, and D. E. Willard. 2019b. Nocturnal flight-calling behaviour predicts vulnerability to artificial light in migratory birds. Proceedings of the Royal Society B 286:20190364.

Zera, A. J., and L. G. Harshman. 2001. The physiology of life history trade-offs in animals. Annual Review of Ecological Systems 32:95–126.

Zúñiga, D., Y. Gager, H. Kokko, A. M. Fudickar, A. Schmidt, B. Neaf-Daenzer, M. Wikelski, et al. 2017. Migration confers winter survival benefits in a partially migratory songbird. eLife 6.

Zurell, D., L. Gallien, C. H. Graham, and N. E. Zimmermann. 2018. Do long-distance migratory birds track their niche through seasons? Journal of Biogeography 45:1459–1468.

